# High-quality thermodynamic data on the stability changes of proteins upon single-site mutations

**DOI:** 10.1101/036301

**Authors:** Fabrizio Pucci, Raphaël Bourgeas, Marianne Rooman

## Abstract

We have set up and manually curated a dataset containing experimental information on the impact of amino acid substitutions in a protein on its thermal stability. It consists of a repository of experimentally measured melting temperatures (*T_m_*) and their changes upon point mutations (Δ*T_m_*) for proteins having a well-resolved X-ray structure. This high-quality dataset is designed for being used for the training or benchmarking of in silico thermal stability prediction methods. It also reports other experimentally measured thermodynamic quantities when available, *i.e*. the folding enthalpy (Δ*H*) and heat capacity (Δ*C_P_*) of the wild type proteins and their changes upon mutations (ΔΔ*H* and ΔΔ*C_P_*), as well as the change in folding free energy (ΔΔ*G*) at a reference temperature. These data are analyzed in view of improving our insights into the correlation between thermal and thermodynamic stabilities, the asymmetry between the number of stabilizing and destabilizing mutations, and the difference in stabilization potential of thermostable versus mesostable proteins.

## I. Introduction

The availability of a complete and well-curated dataset for training and testing purposes is a basic prerequisite for the development of any knowledge-based bioinformatics prediction tool. Here we present a repository containing thermal and thermodynamic stability data on experimentally characterized single-site protein mutants for which an X-ray structure is available, which we have set up by screening the literature and freely accessible databases. This dataset is meant to be as complete as possible, and to contain as much as possible noise- and error-free data. The amount and quality of the data used to set up a predictor are indeed two fundamental requirements for getting reliable predictions.

We have used this dataset to design and test our method for predicting the change in melting temperature of proteins upon point mutations [1], and to compare its performance with that of other existing tools. This dataset is intended as a common benchmark for training and validating different in silico tools for protein stability prediction and rational design.

There is an increasing interest for the development of reliable stability predictors that can be used for rationally designing protein mutants with improved properties, and there is thus concomitantly an increasing need for high-quality and easily accessible datasets. Indeed, the design of new enzymes and other proteins that remain stable and active in unusual environments or at temperatures that differ from their physiological temperatures would allow the optimization of a wide series of biotechnological processes in many sectors such as agro-food, biopharmaceuticals and environment [2, 3].

Another asset of this dataset is that it can serve as a basis for large-scale analyses in view of improving our understanding of the factors that modulate the thermal resistance and other thermodynamic properties of proteins and their variation upon mutation. As a matter en fact, we performed in this paper some analyses that yield some interesting insights.

## II. Brief Theoretical Review of Protein Stability

The protein folding transition is thermodynamically characterized by a change in free energy, enthalpy, entropy and heat capacity. Assuming protein folding to be a two-state transition and the change in heat capacity Δ*C_p_* to be temperature independent, the folding free energy Δ*G(T)* = *G^folded^(T)* – *G^unfolded^(T)* can be written as

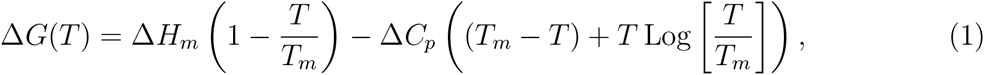

where *T_m_* is the melting temperature and Δ*H_m_* the enthalpic change measured at *T_m_*. Note that with these conventions, Δ*H_m_* and Δ*C_p_* are negative; the folding free energy Δ*G(T)* is negative under physiological conditions and is positive for temperatures above *T_m_*. The protein stability curve has thus an inverse bell shape, as shown in Figure 1.

**FIG. 1:**
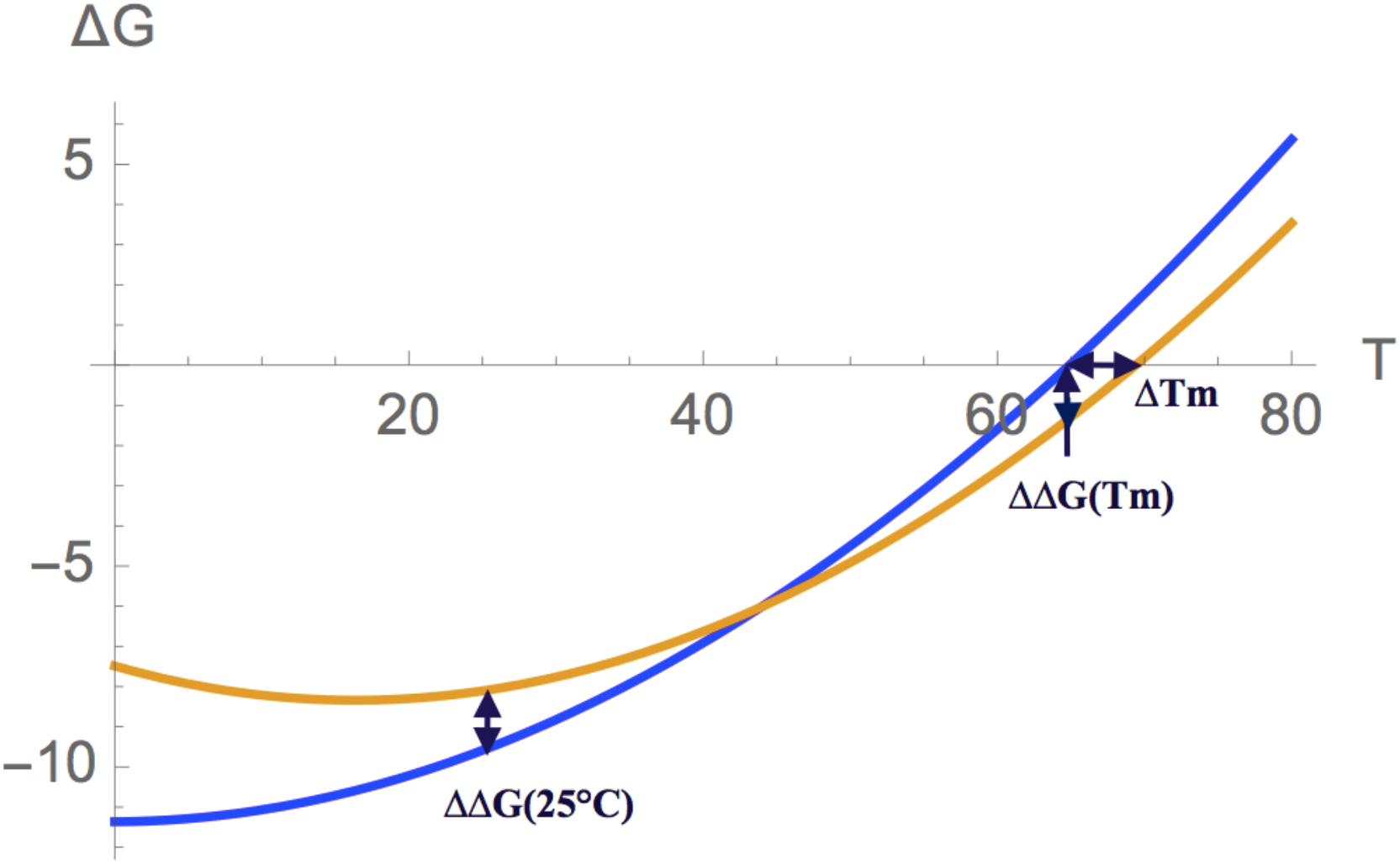
Folding free energy in kcal/mol as a function of the temperature for human lysozyme (PDB code 1LZ1) (blue curve) and its mutant R21A (green curve). While there is an anticorrelation between Δ*T_m_* and ΔΔ*G(T_r_)* at *T_r_* = *T_m_*, there is a correlation between them at *T_r_* = 25°C. This is an example of mutation that thermally stabilizes the protein and thermodynamically destabilizes it.

When one or several residues of the wild type protein are substituted, the change in protein stability due to the mutation can be characterized by a temperature descriptor, namely the change in melting temperature upon mutation:

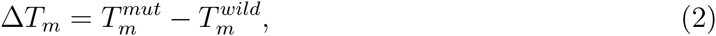

which measures the change in thermal stability. It can also be characterized by a free energy descriptor, *i.e*. the change in folding free energy a the reference temperature *T_r_* (usually chosen to be the room temperature *T_r_* = 298 K):

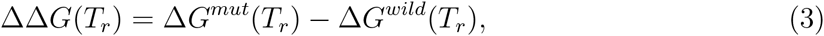

which measures the change in so-called thermodynamic stability. Note that with these conventions, thermally stabilizing mutations have positive Δ*T_m_*-values, and thermodynamically stabilizing mutations have negative ΔΔ*G*(298*K*) values.

The dataset presented in this paper has been created in view of developing a predictor of protein thermal stability changes upon point mutations [1]. It thus contains all the point mutations with experimentally measured Δ*T_m_* values that we have collected from the literature and satisfy certain criteria. The corresponding values of ΔΔ*G(T_r_)* and of the other thermodynamic quantities appearing in equation (1) are known only for a subset of the entries.

In order to understand the precise relation between thermal and thermodynamic stability changes, one needs to have independent experimental determinations of their respective descriptors Δ*T_m_* and ΔΔ*G(T_r_)*, or to know the values of all the thermodynamic quantities that appear in equation (1), *i.e*. *T_m_*, Δ*H_m_*, and Δ*C_P_*, for both the wild type and mutant proteins. Unfortunately all these informations are not always available or not sufficiently accurately measured. Under the assumption that the mutated protein is a perturbation of the wild type, some approximations can be made; for example it is quite reasonable to consider that the parameter

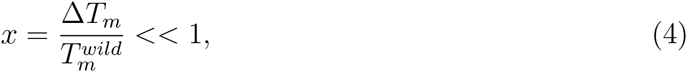

with the temperature expressed in Kelvin, is small and thus that an expansion of equation (3) in powers of *x* can be performed. This yields:

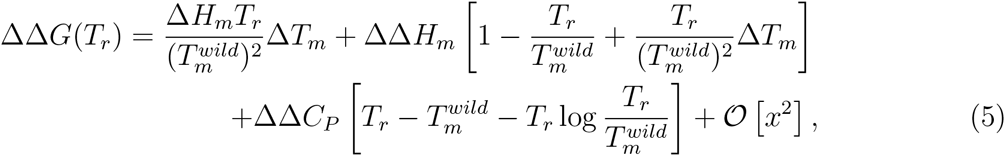

where 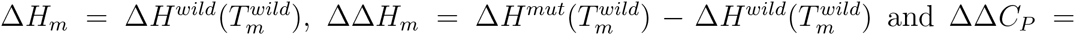 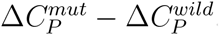. As seen from this equation, the correlation between ΔΔ*G(T_r_)* and Δ*T_m_* generically depends on an intricate combination of variations of thermodynamic quantities. If we assume ΔΔ*H_m_* ⋍ 0 and ΔΔ*C_p_* ⋍ 0, equation (5) reduces to:

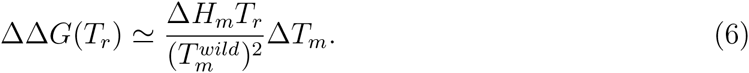

Under this (strong) assumption, we find thus a linear relation between ΔΔ*G(T_r_)* and Δ*T_m_*; the proportionality coefficient is however protein-dependent. Note that Δ*H_m_* is negative with our conventions, and that ΔΔ*G(T_r_)* and Δ*T_m_* are thus anticorrelated.

On the other hand, at the reference temperature 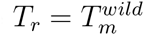, equation (5) simplifies to:

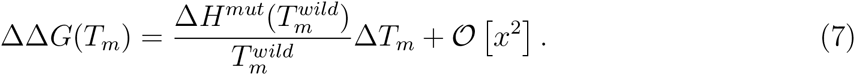

The proportionality assumption between ΔΔ*G(T)* and Δ*T_m_* is thus valid at *T_m_*. If moreover we assume ΔΔ*H_m_* ⋍ 0, this equation becomes the Becktel-Schellman formula [4]:

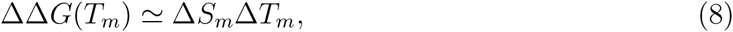

where 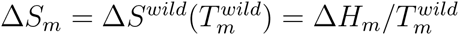 is the entropic contribution at 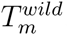.

## III. Methods

### A. Dataset design

We started collecting the mutations with experimentally measured Δ*T_m_* values from the ProTherm database [5], and searched for additional entries by literature screening. Each entry (including those from ProTherm) was manually and carefully checked from the original literature to remove imprecisions and errors. We selected the mutations that satisfy the following criteria:

- Only single point mutations were included.
- Only mutations in proteins, whose three dimensional (3D) structures were experimentally solved by X-ray crystallography with a resolution of at most 2.5 Å, were considered.
- Only mutations that were experimentally characterized in monomeric proteins were taken into account, irrespective of the oligomeric state of the biological unit; this ensures that the measured *T_m_* corresponds to the (un)folding transition and not to a change in quaternary state.
- Only wild-type and mutant proteins that are described in the reference articles as undergoing a two-state (un)folding transition were included.
- Destabilizing or stabilizing mutations by more than 20 °C were overlooked, as they are likely to induce important structural modifications.

When several experimental Δ*T_m_* values were found in the literature for the same mutation, we chose the one measured at pH closest to seven and with the lowest concentration of additives; if more than one measurement in the same conditions was available, the average Δ*T_m_* was taken.

In addition to the change in melting temperature upon mutation, other thermodynamic quantities associated to the mutation are reported in the dataset when available. These are the Δ*C_P_* of the wild type protein and its change upon mutation ΔΔ*C_P_*, ΔΔ*G(T_r_)* and the reference temperature *T_r_* at which the measurement was performed, the Δ*H_m_* of the wild type and

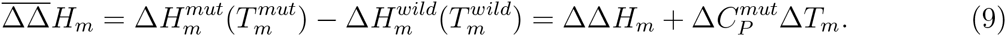

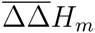 refers to a quantity that is slightly different from ΔΔ*H_m_* appearing in equation (5): the former is computed at different *T_m_* values whereas the latter is computed at 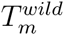; the difference is proportional to Δ*T_m_*. We report in the dataset 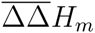 rather than ΔΔ*H_m_* as these are the measured quantities. Note that for a given mutation these different quantities are not always measured in exactly the same experimentally conditions than the corresponding Δ*T_m_* values.

When available, the ΔΔ*G*’s indicated in the dataset are the values that are measured by monitoring the (un)folding transition through chemical (de)naturation using urea or guanidinium chloride (GdmCl); the temperature at which the experiments were performed is also reported. If such data are not available but all the thermodynamic quantities in equation (1) are known for the wild type and the mutant proteins, they are used to evaluate ΔΔ*G* at 25°C. Otherwise, approximations were made to evaluate ΔΔ*G*, and the corresponding entries in the dataset are labeled by a subscript. The ΔΔ*G* values obtained with the approximation consisting in considering ΔΔ*C_P_* ⋍ 0 are indicated with a subscript (*b*); the temperature at which they were estimated is equal to 25°C. When the stronger approximation consisting of supposing also ΔΔ*H_m_* ⋍ 0 (see equation (6)) is used to derive the value of ΔΔ*G* from Δ*T_m_*, we mark it with the subscript (*a*); the temperature at which this quantity is given is equal to 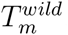. For the few entries whose values are labeled with a subscript (*c*), the ΔΔ*G*(25°C) values are computed from Δ*T_m_* using an empirical correlation between the two quantities computed on a subset of mutants of the same wild type protein [6]. Finally the ΔΔ*G* values at *T_m_* that are derived from the approximation (see ref. [7, 8]) ΔΔ*G(T_m_)* = 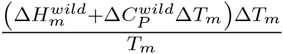 are labeled with the subscript (*d*).

The experimental techniques used for measuring the protein melting temperatures and other thermodynamic quantities are indicated in the dataset. These are differential scanning calorimetry (DSC), circular dichroism (CD), absorbance (Abs), and fluorescence.

The Protein DataBank (PDB) code [9] of the best resolved 3D X-ray structure of each wild type protein is specified in the dataset. For a few entries, the PDB code is labeled with a subscript. This means that the wild type structure of the protein whose Δ*T_m_* was measured was unavailable, and that the structure of an almost identical protein was used instead, under the assumption that the impact of the modification on the structure is negligible. In particular, the 1bni_*h*102*a*_ code means that the structure is obtained from the PDB structure 1bni with the His residue at position 102 manually substituted into an Ala. The same procedure is used for the PDB structures 1ycc_*c*102*a*_, 1urp_*l*265*c*_ and 5pti_*m*52*l*_. The other PDB codes with subscripts, *i.e* 1tpk_*r*_, 1yu5_*d*1_ and 1yu5_*d*2_, refer to experimentally characterized proteins whose sequences have been manually truncated by a few residues compared to the original PDB structure. Note that we checked that the mutations or truncated residues in these pseudo-wild type proteins are all distant from the mutations whose Δ*T_m_* was measured, so that they may be assumed as not interfering.

### B. Data records

The dataset contains experimental information on 1,626 point mutations that have been introduced in about 93 proteins. This data was collected by screening the literature and databases, and carefully checked on the basis of the original articles. For each mutation, the following informations are reported:

- The PDB [9] code of the 3D structure of the wild type protein (Column II).
- The chain name, residue number and residue name of the wild type and mutant amino acids (Columns III-VI).
- The experimental value of the change in melting temperature upon mutation (Δ*T_m_)* using the convention of equation (2) (Column VII).
- The experimentally measured melting temperature (*T_m_*) and the number of residues (*N_r_*) of the wild type protein (Columns VIII and XIV, respectively).
- The experimental values of 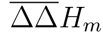, which is the change in calorimetric enthalpy upon mutation measured at the mutant and wild type melting temperatures, respectively, as defined in equation (9) (Column IX), and of the Δ*H_m_* of the wild type protein at 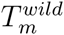 (Column X), when available. The subscript (*e*) means that the reported values correspond to the van’t Hoff enthalpy instead of the calorimetric enthalpy.
- The experimentally measured values of ΔΔ*C_P_* (Column XI) and the 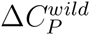 of the wild type protein (Column XII), when available.
- The values of ΔΔ*G(T_r_)*, using the conventions of equation (3) (Column XIII), and the reference temperature *T_r_* (in degrees Celsius) at which they were measured or derived (Column XIV). Entries without superscripts are experimental or calculated from other measured thermodynamic quantities, whereas entries with superscripts (a), (b), (c) or (d) were obtained using different levels of approximations, as explained in the previous section.
- The resolution of the X-ray structure (in Å) (Column XVI).
- The name of the protein and its host organism (Columns XVII-XVIII)
- The bibliographic references (Column XIX).
- The pH and the experimental technique used for measuring Δ*T_m_* (Columns XX-XXI).

## IV. Results

We investigated some biophysical properties of the data reported in our dataset. First of all, the Δ*T_m_* distribution obtained from all the entries is dominated by destabilizing mutations, as shown in Figure 2. The average *T_m_*-value, 〈Δ*Τ_η_*〉, is indeed equal to −2.7°C, while the standard deviation and the kurtosis of the distribution are equal to 5.3°C and 4.0°C, respectively. The large majority of point mutations (about 70%) are thus destabilizing. Although this Δ*T_m_* distribution is not built from the ensemble of possible mutations but rather from the subset of experimentally characterized mutations, we may nevertheless assume that it represents well the actual Δ*T_m_* distribution of all possible mutations. The relative abundance of destabilizing mutations with respect to stabilizing ones can be interpreted as being due to the evolutionary force that tends to optimize the proteins for stability and thus to minimize the deleterious impact of random mutations. It must nevertheless be emphasized that all proteins are left with stability weaknesses [10] or frustrations [11], in particular in functional regions.

**FIG. 2:**
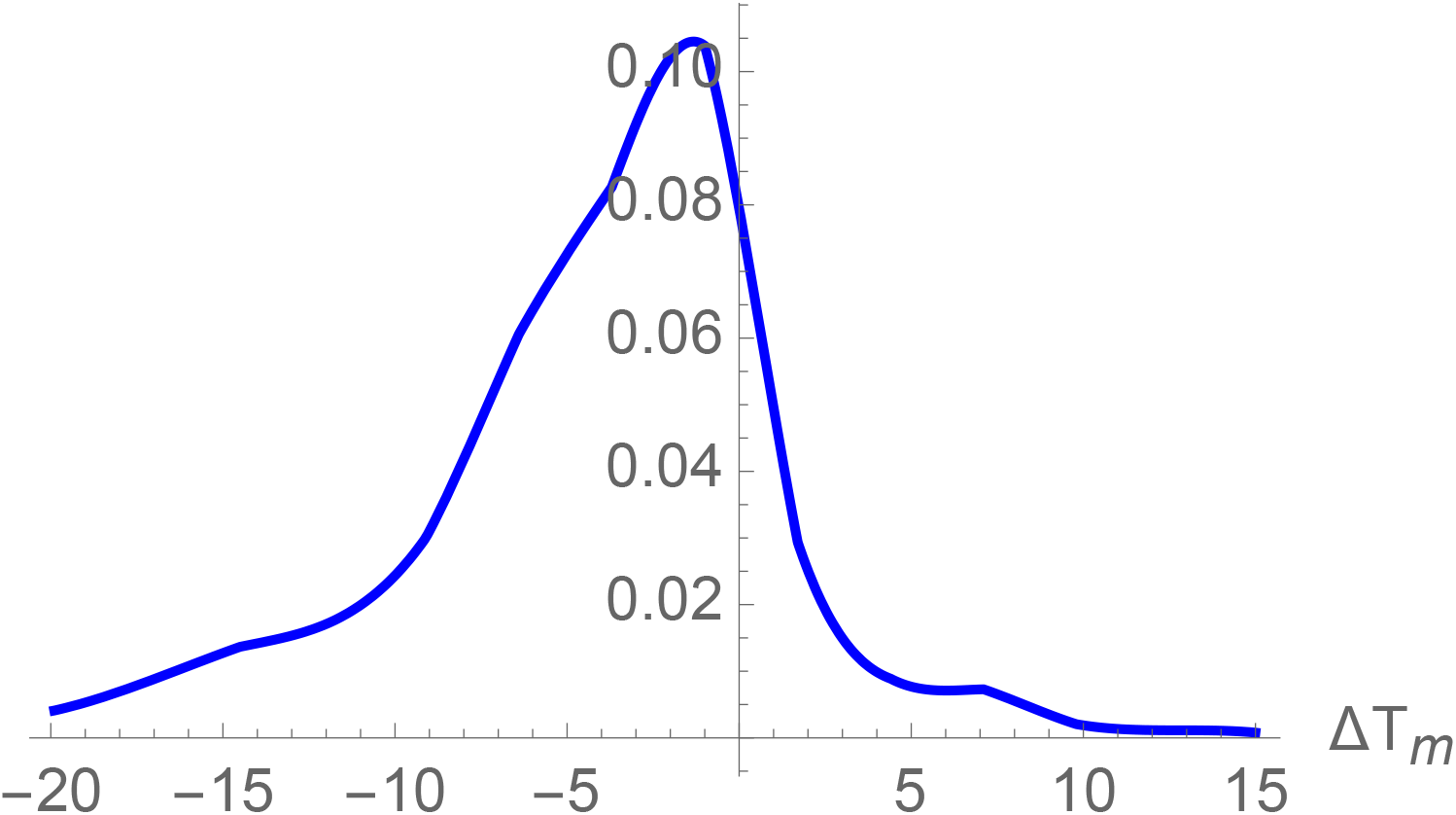
Normalized Δ*T_m_* distribution for the 1,626 mutations collected in the dataset.

The solvent accessibility of the mutated residues is an important feature that modulates the average stability changes. It is defined as the ratio between the solvent accessible surface of a residue X in a given structure and in the extended tripeptide Gly-X-Gly conformation, and has been computed using an in-house program [12]. In Figures 3a-c, we show the experimental Δ*T_m_* distribution as a function of the solvent accessibility of the mutated residues; three solvent accessibility (Acc) ranges are considered: Acc < 15% (core), 15% < Acc < 50% (partially buried) and Acc > 50% (surface). The three distributions were found to be significantly different according to the 2-sample Kolmogorov-Smirnov (K-S) test (P-value < 10^—3^). The mean 〈Δ*T_m_*〉 values of the distributions are equal to −4.3°C, −1.6°C and −1.1°C for the core, partially buried and surface mutations, respectively. As expected, the mutations in the core are on the average more destabilizing than those at the surface since core residues play a stronger role in the structural stability than surface residues, which also contribute to stability but to a lesser extent.

**FIG. 3:**
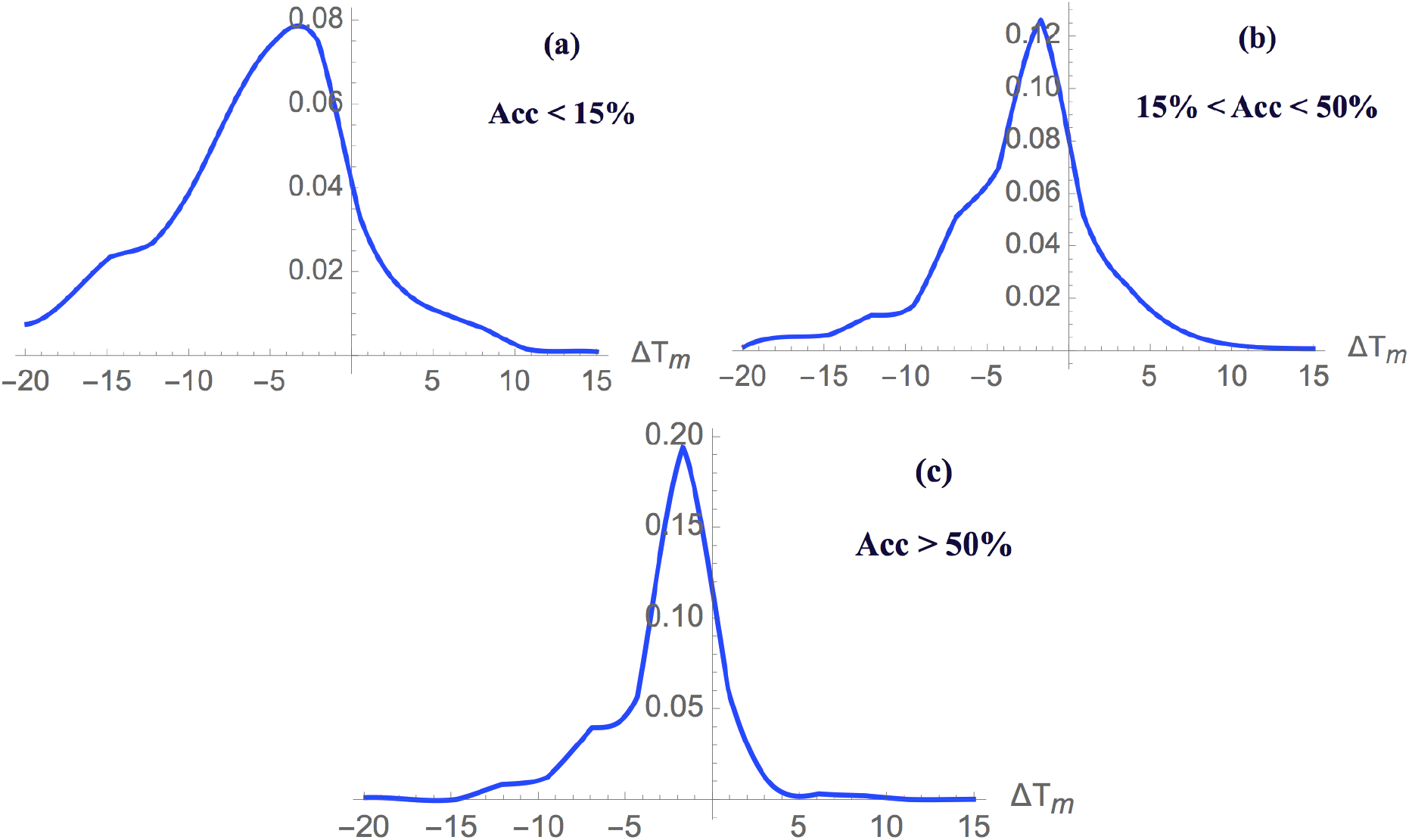
Normalized Δ*T_m_* distribution for: (a) 734 mutations in the protein core (Acc<15%), (b) 513 mutations of partially buried residues (15%<Acc<50%) and (c) 379 mutations at the protein surface (Acc>50%)

It is also informative to analyze the relation between Δ*T_m_* and *T_m_*. Indeed, it could be argued that it is "easier" to destabilize thermostable proteins or equivalently, to stabilize mesostable proteins. To check this hypothesis, we computed 〈Δ*T_m_*〉 separately for the mutations introduced in thermostable proteins defined here as having a melting temperature higher than 65°C and those introduced in mesostable proteins with *T_m_* < 65°C. We obtain a value of 〈Δ*T_m_*〉 = –3.6°C for thermostable proteins and 〈Δ*T_m_*〉 = –2.1°C for mesostable proteins. On the average, mutations in thermostable proteins are thus more destabilizing than in mesostable proteins. Moreover, the normalized Δ*T_m_* distributions for mesostable and thermostable proteins are shown in Figure 4. They are statistically different according to the K-S test with a P-value < 10^−4^, and the former appears to be shifted towards stabilizing mutations compared to the latter. This interesting result supports the view that the fraction of stabilizing mutations is larger in mesostable proteins than in thermostable proteins, and thus that the former are easier to stabilize than the latter, in agreement with the starting hypothesis.

**FIG. 4:**
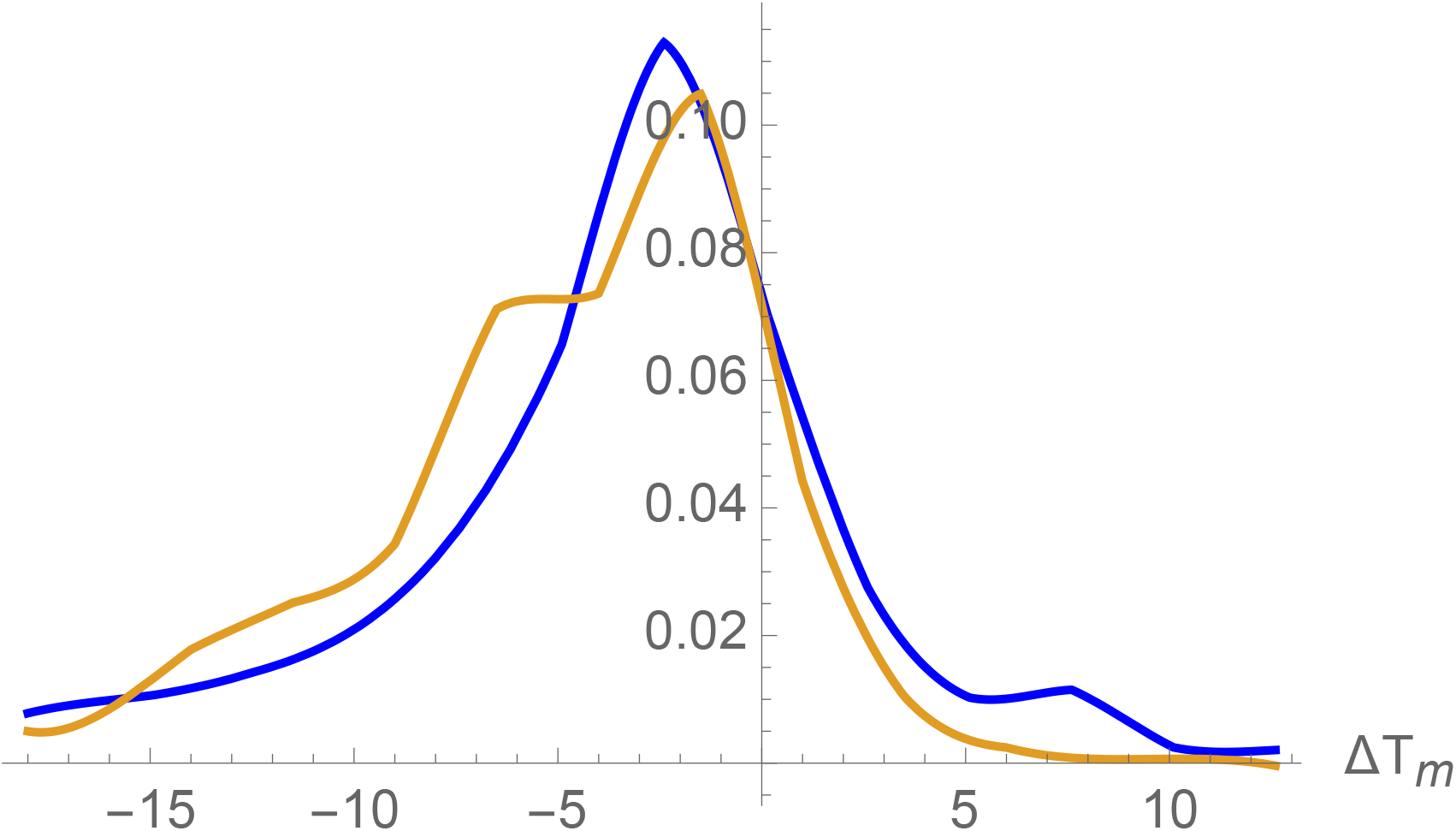
Normalized Δ*T_m_* distributions for mesostable (blue) and thermostable proteins (orange).

We also analyzed the other thermodynamic quantities reported in our dataset and first of all, the change in folding enthalpy upon point mutations. The normalized 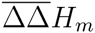 distribution (see equation (9)) is plotted in Figure 5a; its mean value is 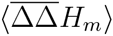 = 7.3 kcal/mol for the set of 993 entries for which this quantity has been measured experimentally. Hence, the mutations are on the average enthalpically destabilizing at *T_m_*. Figure 5b shows the normalized distribution of the change in entropy upon mutation 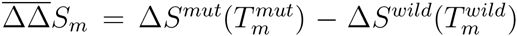. The mean value of the distribution is found to be 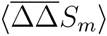 = 0.04 kcal/(mol K).

Finally, we plotted the normalized distribution of ΔΔ*C_P_* and ΔΔ*G* in Figures 5c-d. The mean values of these two distributions are positive: 〈ΔΔ*C_p_*〉 = 0.08 kcal/(mol K) for the set of 250 entries for which this value has been measured, and 〈ΔΔ*G*〉 =0.89 kcal/mol for 1,147 entries. Hence, the majority of mutant proteins have a less negative Δ*C_P_* and a less negative Δ*G* than wild type proteins; the mutant proteins are thus on the average less thermodynamically stable than the wild type. Note that the asymmetric nature of the ΔΔ*G* distribution is likely to cause biases in the prediction methods that use these data as learning set [13].

In summary, the large majority of the mutations are thermodynamically destabilizing (as measured by positive ΔΔ*G*) and thermally destabilizing (as measured by negative Δ*T_m_*).

**FIG. 5:**
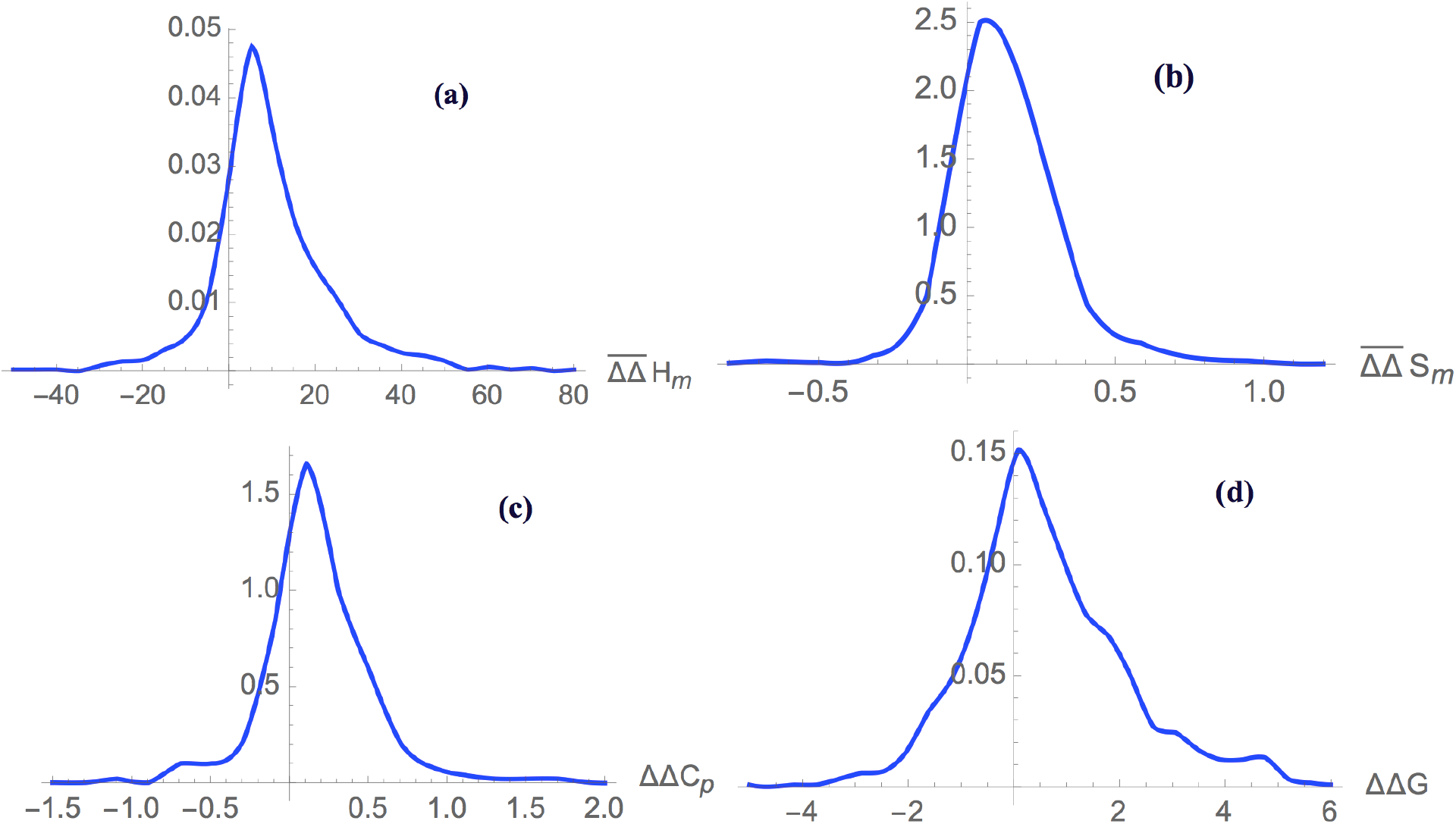
Normalized distributions for: (a) 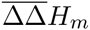 (in kcal/mol) and (b) 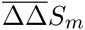 (in kcal/(mol K)) obtained from 993 mutations, (c) ΔΔ*C_p_* (in kcal/(mol K)) obtained from 250 mutations, and (d) ΔΔ*G* (in kcal/mol) obtained from 1,147 mutations.

The next point we investigated is the correlation between the thermodynamic stability descriptor ΔΔ*G(T_r_)* and the thermal stability descriptor Δ*T_m_*. Indeed, these two quantities are often taken as equivalent stability measures even though this assumption is based on an approximation, as shown in equation (6). Nevertheless, this hypothesis seems *a priori* not totally unjustified, as the linear anticorrelation between the two quantities is in general quite good. In our dataset, the Pearson correlation coefficient *r*, computed on the 1,147 mutations for which both ΔΔ*G(T_r_)* and *ΔT_m_* are available independently from the choice of *T_r_* is equal to:

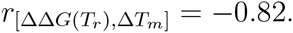

We must however notice that the temperature at which the ΔΔ*G* measurements were performed is not always the same (as described in the "Dataset design" subsection of Methods); it is usually either 25°C or the *T_m_* of the wild type protein.

For the subset of 461 mutations for which ΔΔ*G* has been measured or computed at 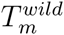, the linear anti-correlation is close to perfect:

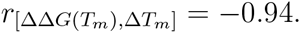

The anticorrelation does not reach −1 since the proportionality coefficient between ΔΔ*G(T_m_)* and Δ*T_m_* is protein-dependent (see equations (6)-(7)). It is illustrated in Figure 6b.

In contrast, for the 449 mutations for which ΔΔ*G*(25°C) has been directly measured or for which all thermodynamic quantities that allow using the full equation (5) have been measured (entries without subscript in the dataset), the anticorrelation is much lower:

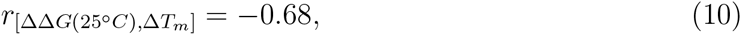

as shown in Figure 6a. We would like to stress the value of this ΔΔ*G*(25°C)-Δ*T_m_* anticorrelation coefficient can be expected to be close to the real one, as it has not been artificially improved by adding computed ΔΔ*G*’s that presuppose this anticorrelation.

For some entries, the two descriptors ΔΔ*G*(25°C) and Δ*T_m_* are correlated rather than anticorrelated. These signal interesting mutations that stabilize the protein thermally while destabilizing it thermodynamically at room temperature, or conversely, destabilize it thermally while stabilizing it thermodynamically. As an example of such an unusual behavior, we plotted in Figure 1 the full protein stability curve of the wild type human lysozyme and of the mutant R21A [14]; these are one of the entries of our dataset.

**FIG. 6:**
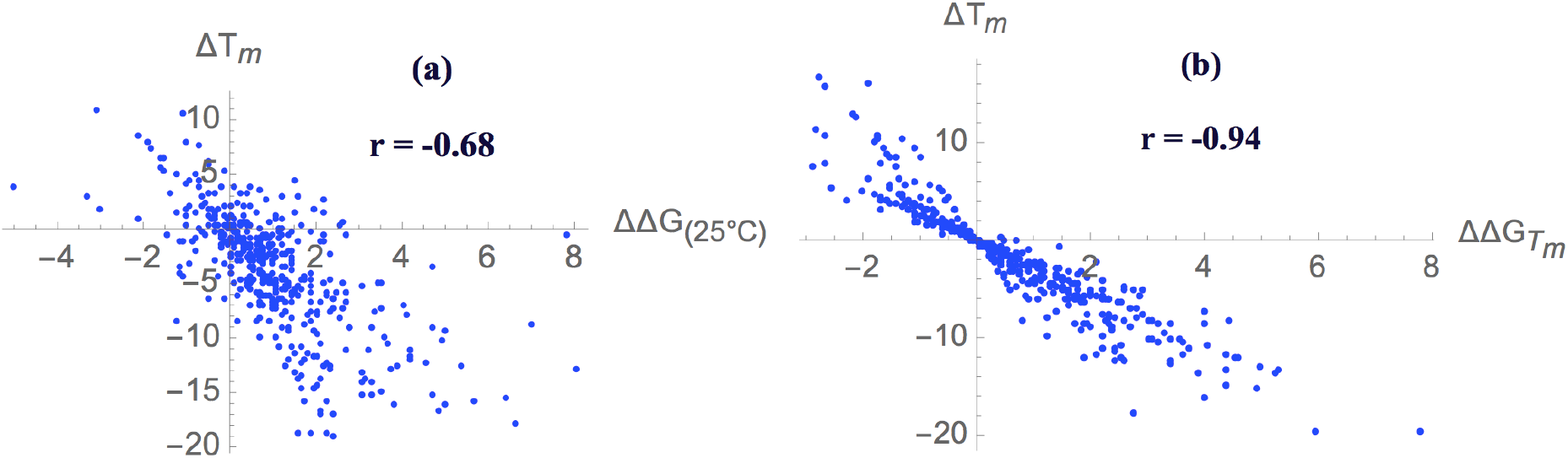
Anticorrelation between the two descriptors of protein stability. (a) Anticorrelation between Δ*T_m_* and ΔΔ*G*(25°C) and (b) anticorrelation between Δ*T_m_* and ΔΔ*G(T_m_)* where *T_m_* is the melting temperature of the wild type protein.

### Sources of experimental errors

The experimental errors on the measured thermodynamic quantities describing the folding transition have to be taken in consideration. The most noisy thermodynamic quantities are Δ*C_P_*. Their error is generally of the order of 10-20%. Sometimes it is of the same order as ΔΔ*C_P_* itself, which makes the numeric evaluation of equation (5) not quite precise, even though the ΔΔ*C_P_* term is subleading compared to the others.

The errors on the two thermodynamic descriptors Δ*H_m_* and *T_m_* are in general less severe, being of the order of a few percents. These should thus not really affect the results obtained in this analysis.

Another source of error comes from the fact that the experiments are often performed in different environmental conditions in terms of pH, buffer type, ionic concentration and additives. Such errors are non negligible, even if their effect can be expected to be less important for the variation of the thermal characteristics upon mutations compared to that of the thermal characteristics themselves. Moreover, to decrease this effect, we have collected data that are as much as possible uniform in terms of environmental variables, as explained in the section "Dataset design". Note that the size of this type of error is difficult to quantify in general.

### Usage notes

The dataset that we have constructed is available as a pdf file in attachment to this paper and can be downloaded as a text file at the address http://babylone.ulb.ac.be.

## Acknowledgments

We acknowledge support from an FRFC grant from the Belgian Fund for Scientific Research (FNRS). RB is a Postdoctoral Fellow, FP a Postdoctoral Researcher and MR a Research Director at the FNRS.

## Data Citations

Bibliographic information for the data records described in the manuscript.

